# Multi-ancestry Genome- and Phenome-wide Association Studies of Diverticular Disease in Electronic Health Records with Natural Language Processing enriched phenotype algorithm

**DOI:** 10.1101/2020.06.08.138735

**Authors:** Yoonjung Yoonie Joo, Jennifer A Pacheco, William K Thompson, Laura J Rasmussen-Torvik, Luke V Rasmussen, Frederick TJ Lin, Mariza de Andrade, Kenneth M Borthwick, Erwin Bottinger, Andrew Cagan, David S Carrell, Joshua C Denny, Stephen B Ellis, Omri Gottesman, James G Linneman, Jyotishman Pathak, Peggy L Peissig, Ning Shang, Gerard Tromp, Annapoorani Veerappan, Maureen E Smith, Rex L Chisholm, Andrew Gawron, Abel N Kho, M Geoffrey Hayes

## Abstract

**Background and aims:** Diverticular disease is among the most prevalent conditions encountered by gastroenterologists, affecting ∼50% of Americans before the age of 60. Our aim was to identify genetic risk variants and clinical phenotypes associated with diverticular disease, utilizing the electronic health record (EHR) with Natural Language Processing (NLP).

**Methods:** We developed a NLP-enriched phenotype algorithm that incorporated colonoscopy or abdominal imaging reports to accurately identify patients with diverticulosis and diverticulitis from multicenter EHRs. We performed genome-wide association studies (GWAS) of diverticular disease in European, African and multi-ancestry participants, followed by phenome-wide association studies (PheWAS) of the risk variants to identify their potential comorbid/pleiotropic effects in the clinical phenome. For more in-depth investigation of associated clinical phenotypes, we also performed PheWAS with the previously reported 52 GWAS susceptibility variants for diverticular disease.

**Results:** Ancestry-stratified GWAS analyses confirmed the well-established associations between *ARHGAP15* loci with diverticular disease in European cohorts, and found similar positive effect sizes in African cohorts but with non-significant p-values. With overall intensified GWAS signals in diverticulitis patients compared to diverticulosis patients, we found substantial genetic correlations between diverticulosis and diverticulitis, up to 0.997 in European ancestry. PheWAS analyses identified associations between the diverticular disease GWAS variants and circulatory system, genitourinary, and neoplastic EHR phenotypes.

**Conclusion:** Our multiancestry GWAS-PheWAS study demonstrated an effective use of multidimensional EHR information in disease case/control classification with NLP for more comprehensive and scalable phenotyping, and implementation of an integrative analytical pipeline to facilitate etiological investigation of a disease from a clinical perspective.

## Introduction

Diverticular disease is the most common morphological defects of the intestinal tract and the fifth most important gastrointestinal (GI) disorder in terms of medical cost as high as >$5.4 billion in the United States^1-3^. Diverticular disease usually indicates asymptomatic diverticulosis (the mere presence of diverticula, a pouch-like protrusion in the colonic wall), but also includes diverticulitis (an acute or chronic inflammation of diverticula) and its clinical complications^4^. Diverticulitis occurs in approximately 4% to 15% of patients with diverticula with a high reoccurrence rate, which is associated with fever, abdominal pain, leukocytosis and potential life-threatening peritonitis^4-8^.

The disease is highly prevalent in Western countries that have achieved a high degree of industrialization and urbanization^9^. North America has the highest prevalence of diverticular disease, where it is estimated to be 5-10% of the population younger than 40, ∼33% of the population older than 45, and up to 67% of the population older than 65^7, 10^. The high prevalence in Western countries is in contrast to that in the countries that do not follow Western lifestyles, e.g. the overall prevalence in Asia and Africa is estimated to be <0.5 to 25%^5 9^. Despite this geographic variation, virtually all countries worldwide are observing an increasing burden of diverticular disease irrespective of their economic developmental or demographical variability. In Finland, the incidence of diverticular disease has risen by 50% in the last two decades^11^. The US hospitalization rate for acute diverticulitis has increased 26% between 1998 to 2005^12^, and a similar pattern is observed in Japan^13^, Canada^14^, England^15^, Singapore^16^, Nigeria^17^, and South Africa^18^. Dietary intake of low fiber, processed foods, and red meats, have been implicated as potential causes of diverticular disease^9, 19, 20^, but is controversial^21, 22^.

As with most conditions, current evidence supports a complex interplay of both environmental and genetic contributions. Twin studies reveal that genetic heritability of diverticular disease is estimated to be up to 53% (95% Confidence Interval (CI), 45-61%)^6^. To date, three GWAS have identified 52 genetic susceptibility loci associated with diverticular disease^23-25^.

A significant challenge to etiologic investigation is that approximately 75% to 90% of the diverticulosis patients stay asymptomatic until presenting with diverticulitis^26^, making it difficult to self-identify or detect the disorder in a clinical setting. In the acute setting a computed tomography (CT) imaging of the abdomen is most often used in the evaluation of diverticulitis, but it may not be completely diagnostic in cases of early or mild diverticulitis^27^. Currently, the definitive ascertainment of the presence or absence of diverticular disease depends on undergoing a colonoscopy^5, 27, 28^, but this requirement suffers from incomplete compliance of patients with current screening guidelines^29^.

To address limitations with previous research, we conducted a GWAS in the electronic Medical Records and Genomics (eMERGE) network, a collaborative consortium of multiple medical institutions in the United States with the capacity and patient consent to link genomic data of patients with EHR data^30, 31^. For our study, eMERGE sites collected clinical diagnosis codes, demographic information, colonoscopy and abdominal imaging reports from the EHR, which was combined with genetic information for deeper etiologic investigation of diverticular disease.

In this study, we developed a phenotype algorithm that incorporated Natural Language Processing (NLP) to identify the presence or absence of diverticulosis or diverticulitis from EHRs with high accuracy. We aimed to present a scalable framework to discover clinical pleiotropy of common genetic risk variants through EHR-powered GWAS and phenome-wide association studies (PheWAS).

## Methods

### NLP-enriched phenotype algorithm for diverticular disease

Genome-wide genotype data of 38,827 individuals from 9 biobanks and phenotype data including their demographic, clinical diagnosis, colonoscopy or abdominal imaging reports of 99,185 individuals were collected from 12 biobanks in the eMERGE consortium^31^. The details of genotyping, imputation, and quality control processes are explained in **Supplementary material**.

We developed a phenotype algorithm with two variants: each of which incorporated structured and NLP data while accounting for data availability at each implementing site. The primarily NLP-driven variant (supplemented with diagnostic and procedure codes) used colonoscopy or abdominal imaging reports to identify diverticular disease **(Figure 1a)**. This was implemented at five eMERGE sites: Northwestern University (NU), Vanderbilt University (VU), Geisinger, Kaiser Permanente Washingon / University of Washington (KPWA/UW), and Mayo Clinic. Two sites (Marshfield, Mount Sinai) had a limited subset of imaging reports available. We developed a second variant of the algorithm to select diverticulosis cases, with or without diverticulitis, primarily using ICD-9 diagnosis codes that started with 562 (‘Diverticulosis and diverticulitis’ category’), assigned within 7 days after a colonoscopy or abdominal imaging **(Figure 1b)**. This structured data was supplemented with NLP when reports were available. Additional criteria to define ‘diverticulosis’ and ‘diverticulitis’ are elucidated in **Supplementary material**.

**Figure 1.**
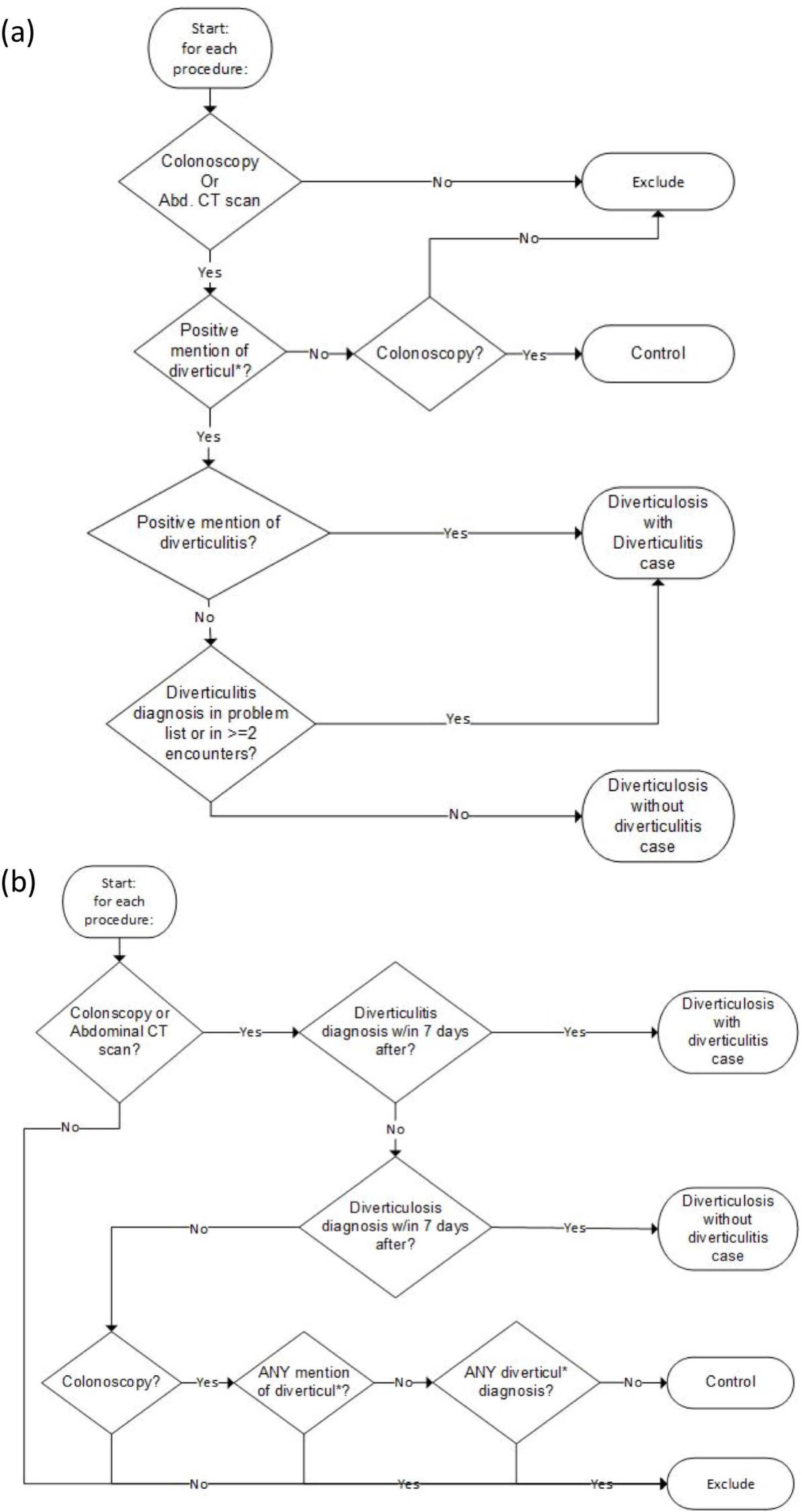
Natural language processing (NLP)-enriched phenotyping algorithms for diverticular disease cases and controls. (a) The NLP-driven phenotype algorithm used in five medical institutions in the eMERGE network (NU, VU, Geisinger, KPWA/UW, Mayo clinic). (b) The structured data-driven phenotype algorithm that was used in two eMERGE sites (Marshfield, Mount Sinai).

Four sites (NU, VU, Geisinger, Marshfield) validated algorithm performance by a standardized chart review of randomly selected patients’ charts. Trained clinicians and chart reviewers reviewed a total of 364 individuals’ records to assess the positive predictive value (PPV) of our developed algorithms, using established guidelines^32^.

### Genome-wide Association Tests

Multi-ancestral (MA) GWAS was conducted on the identified subjects from the 9 sites who implemented our phenotyping algorithm **(Table 1)**. We used logistic regression (PLINK v. 1.9^33^), adjusting for sex, age at colonoscopy, study site, and the first 10 principal components of ancestry. To test for associations with diverticulosis, we compared the patients with diverticulosis, either with or without diverticulitis, to the healthy control patients without any evidence of diverticulosis or diverticulitis. To test for associations with diverticulitis, we excluded any diverticulosis patients without diverticulitis records, and compared the patients with diverticulitis (presenting both diverticulosis and diverticulitis) to the healthy control patients. Similar GWAS were repeated in European ancestry (EA) and African ancestry (AA) participants separately, which are the two largest ancestral groups available. We annotated the significant GWAS loci with eQTL, deleteriousness score (CADD score^34^), and potential regulatory functions (RegulomeDB score^35^) using the GTEx v7 database. A subsequent conditional analysis was performed within a window of ±1Mb of the genome-wide significant GWAS variants using genome-wide complex trait analysis (GCTA) v.1.26^36^.

**Table 1:** Demographic characteristics of the patients by each eMERGE site. Patients with diverticulitis are a subset of the people with diverticulosis.

| Site* | Subjects (N) | Diverticulosis Cases | Diverticulitis Cases | Healthy Controls** | Average Age (mean +/- SD) | Average BMI (mean +/- SD) | Sex (Female) | Race (EA)*** | Race (AA)*** |
| --- | --- | --- | --- | --- | --- | --- | --- | --- | --- |
| All | 21777 | 12577(57.8%) | 1265(10.1%) | 9200(42.2%) | 62.5(+/-12.2) | 29.4(+/-6.8) | 11891(54.6%) | 19211(88.2%) | 2322(10.7%) |
| Columbia University | 523 | 39(7.5%) | 39(100.0%) | 484(92.5%) | 63.0(+/-15.4) | 27.9(+/-10.4) | 283(54.1%) | 311(59.5%) | 168(32.1%) |
| Kaiser Permanente Washington/University of Washington | 862 | 448(52.0%) | 120(26.8%) | 414(48.0%) | 74.4(+/-9.8) | 27.3(+/-5.4) | 499(57.9%) | 789(91.5%) | 33(3.8%) |
| Geisinger | 1603 | 1093(68.2%) | 170(15.6%) | 510(31.8%) | 65.6(+/-12.4) | 30.7(+/-7.4) | 596(37.2%) | 1595(99.5%) | 6(0.4%) |
| Harvard | 1651 | 884(53.5%) | 72(8.1%) | 767(46.5%) | 57.4(+/-12.7) | 28.8(+/-6.4) | 943(57.1%) | 1535(93.0%) | 90(5.5%) |
| Marshfield | 3324 | 2214(66.6%) | 266(12.0%) | 1110(33.4%) | 62.8(+/-10.0) | 29.5(+/-5.7) | 1971(59.3%) | 3306(99.5%) | 2(0.1%) |
| Mayo | 5417 | 3275(60.5%) | 251(7.7%) | 2142(39.5%) | 65.1(+/-10.9) | 29.3(+/-6.4) | 2509(46.3%) | 5379(99.3%) | 12(0.2%) |
| Mount Sinai | 1133 | 416(36.7%) | 121(29.1%) | 717(63.3%) | 59.0(+/-10.2) | 30.5(+/-7.4) | 694(61.3%) | 339(29.9%) | 773(68.2%) |
| NU | 1933 | 993(51.4%) | 77(7.8%) | 940(48.6%) | 57.6(+/-11.4) | 28.8(+/-7.4) | 1496(77.4%) | 1660(85.9%) | 265(13.7%) |
| VU | 5331 | 3215(60.3%) | 149(4.6%) | 2116(39.7%) | 60.8(+/-13.0) | 29.6(+/-7.3) | 2900(54.4%) | 4297(80.6%) | 973(18.3%) |

\*Sites: GHC/UW = Group Health Cooperative/University of Washington, NU = Northwestern University, VU = Vanderbilt University.
\*\* Without diverticulosis or diverticulitis
\*\*\*Race & Ethnicity categories are mutually exclusive: EA = European American, AA = Black or African American; <1% Other race.

### Evaluation of our NLP-enriched phenotype algorithm for diverticular disease

To evaluate, we compared our NLP-enriched phenotype algorithm results against the results of an ICD-based phenotyping method that has been commonly implemented in previous GWASs of diverticular disease^23-25^. Using the phecode map v1.2^37^ for diverticular disease (ICD-9 562), we excluded patients with any related gastrointestinal manifestations such as ‘ulcerative enterocolitis’ (ICD- 9 556), ‘regional enteritis’ (ICD-9 558), ‘volvulus of intestine’ (ICD-9 560.2), etc. for more accurate patient classification. **(Supplementary Table 1)**

### LD score regression

To measure the extent of genetic overlap between diverticulosis and diverticulitis, LD score regression was used to calculate their genetic correlation and SNP-based heritability (due to common variation)^38, 39^. We also calculated the explained heritability by the MA, EA, and AA GWAS results and transformed the estimates into a liability scale, setting population prevalence as 41.7% (MA and EA) and 0.5% (AA) for diverticulosis and 0.02% (MA and EA) and 0.001% (AA) for diverticulitis^6^.

### PheWAS

We conducted PheWAS of independent GWAS-significant SNPs with suggestive threshold (GWAS p-value<1E-06 and LD r^2^<0.1) grouped by ancestry^40^. We retrieved the diagnoses of the 91,166 MA participants, including both ICD-9 and 10 codes, as both were available at the time of analysis. With a minimum of 30 cases per phenotype^40^, logistic regression between the GWAS SNPs and each phecode was performed with the adjustment for the first 10 PCs, and participation sites, through the PheWAS R package^40^. A false discovery rate (FDR) < 0.05 was used for reporting significance.

We also conducted PheWAS of the 52 reported GWAS susceptibility loci from the three existing GWASs of diverticular disease^23-25^. The genomic position of the 52 loci were converted to GrCh37/hg19 (40 loci from Maguire et al^23^, 12 loci from Schafmayer et al^24^), including 3 proxy variants (R^2^ > 0.5) available in our genotype data **(Supplementary Table 5)**.

## Results

### Performance of NLP-enriched phenotype algorithm

As compared against a gold standard of manual clinical chart review, the overall PPV of our phenotype algorithm for diverticulosis cases (with or without diverticulitis) was 0.96, and 0.94 for controls without diverticulosis or diverticulitis. **(Table 2)** We identified 21,777 study participants using the developed algorithm with covariate data. Of these, we identified 12,577 diverticulosis cases with or without diverticulitis, of which 1,265 were diverticulitis cases, and 9,200 controls without diverticulosis or diverticulitis in the entire MA discovery cohort. **(Table 1)**.

**Table 2:** Phenotype algorithm validation and comparison of two phenotyping algorithms for diverticular diseases by site, out of 21,777 subjects with colonoscopy reports.

|  | Diverticulosis Cases Identified |  | Diverticulitis Cases Identified |  | Healthy Controls Identified |  | <u>Evaluation of our algorithm</u> |  |  |
| --- | --- | --- | --- | --- | --- | --- | --- | --- | --- |
|  | NLP-enriched phenotyping algorithm | (Traditional ICD-based phenotyping algorithm)* | NLP-enriched phenotyping algorithm | (Traditional ICD-based phenotyping algorithm) | NLP-enriched phenotyping algorithm | (Traditional ICD-based phenotyping algorithm) | Cases reviewed | Controls Reviewed | PPV** (case/control) |
| **All** | 12,577 | (3,591) | 1,265 | (1,201) | 9,200 | (13,633) | 225 | 139 | 0.96/0.94 |
| **Columbia University** | 39 | (164) | 39 | (40) | 484 | (274) | NA | NA | NA |
| **Kaiser Permanente Washington/University of Washington** | 448 | (227) | 120 | (117) | 414 | (465) | NA | NA | NA |
| **Geisinger** | 1,093 | (401) | 170 | (170) | 510 | (869) | 34 | 33 | 0.97/0.94 |
| **Harvard** | 884 | (482) | 72 | (79) | 767 | (842) | NA | NA | NA |
| **Marshfield** | 2,214 | (655) | 266 | (263) | 1,110 | (1,964) | 50 | 50 | 1.00/1.00 |
| **Mayo** | 3,275 | (695) | 251 | (291) | 2,142 | (3,255) | NA | NA | NA |
| **Mount Sinai** | 416 | (231) | 121 | (80) | 717 | (654) | NA | NA | NA |
| **NU** | 993 | (86) | 77 | (46) | 940 | (1,558) | 91 | 56 | 0.98/0.89 |
| **VU** | 3,215 | (650) | 149 | (115) | 2,116 | (3,752) | 50 | NA | 0.88/NA |

\*This is for comparison purpose. Our main analysis did not utilize the samples identified by this ICD-based phenotyping algorithm.
\*\*PPV = positive predictive value of the phenotyping algorithms overall, and by site, where cases are patients with diverticulosis (either with or without diverticulitis) and controls are patients without diverticulosis (nor diverticulitis), identified by the phenotyping algorithms.

### Evaluation of NLP-enriched phenotyping vs. ICD-based phenotyping

We identified more cases and controls using ICD-based phenotyping, than with NLP-enriched phenotyping, due to the lower availability of report data: 3,313 diverticulitis cases and 45,111 healthy controls with ICD-based phenotyping. However, out of 21,777 subjects with imaging reports data, ICD-based phenotyping identified only 3,591 of them as diverticulosis cases whereas our NLP-enriched algorithm identified 12,577 diverticulosis cases. For diverticulitis, our NLP-enriched algorithm identified 1,265 patients and ICD-based phenotyping identified 1,201 patients (**Table 2)**, and only 87.0% (n=1,101) of case patients were overlapping between these two phenotyping algorithms. Even though the reported PPV of diverticular disease ICD-10 code is as high as 0.98^41^, we find that considerable phenotyping heterogeneity existed without the supporting procedure reports.

### Genetic associations with diverticular disease

The GWAS of diverticular disease in the MA population identified one genome-wide significant locus (**Figure 2**) at 2q22.3 within the *ARHGAP15* gene, which encodes Rho GTPase activating protein 15, and has been consistently reported in previous GWAS of diverticular disease^23-25^. The association patterns between two conditions are largely similar; the diverticulitis GWAS showing more significant and larger ORs compared to the diverticulosis GWAS’s in general (**Table 3**). In the MA GWAS for diverticulosis, one SNP showed eQTL associations with colonic tissues: rs2835676 (*DSCR9* gene) shows strong eQTL association with both transverse and sigmoid colon tissues within the *PIGP* and *TTC3* genes (FDR < 3.90E-13).

**Table 3:** Genetic variants that reach suggestive genome-wide significance (P < 1E-06) with diverticulosis or diverticulitis in MA (multi-ancestry), EA (European ancestry) and AA (African ancestry) participants.

|  | SNP | CH<br>R | POS | Effect<br>Allele | EAF | P-value | OR | SE | Nearest Genes | Function | CADD<br>* | RDB*<br>* |
| --- | --- | --- | --- | --- | --- | --- | --- | --- | --- | --- | --- | --- |
| Diverticulosis<br>MA | **rs6736741** | **2** | **144278534** | **C** | **0.180** | **6.77E-09** | **1.17** | **0.03** | **ARHGAP15:RP11-570L15.2:RP11-570L15.1** | **ncRNA_intronic** | **0.021** | **5** |
|  | rs11200204 | 10 | 123633725 | A | 0.048 | 6.00E-07 | 1.26 | 0.05 | ATE1 | intronic | 0.912 | 6 |
|  | rs2835676 | 21 | 38591311 | T | 0.366 | 7.29E-07 | 0.90 | 0.02 | DSCR9 | ncRNA_intronic | 0.859 | 1f |
| Diverticulitis<br>MA | **rs10928187** | **2** | **144352544** | **G** | **0.225** | **6.63E-12** | **1.42** | **0.05** | **ARHGAP15:RP11-570L15.1** | **ncRNA_intronic** | **0.004** | **7** |
|  | rs56116508 | 4 | 12669861 | C | 0.104 | 1.58E-07 | 1.44 | 0.07 | RP11-352E6.1 | intergenic | 3.290 | 7 |
|  | rs9565028 | 13 | 35704950 | C | 0.415 | 2.53E-07 | 0.79 | 0.05 | NBEA | intronic | 0.173 | 6 |
|  | rs77643000 | 11 | 118749563 | C | 0.131 | 4.23E-07 | 1.38 | 0.06 | CXCR5 | intergenic | 3.446 | 5 |
| Diverticulosis<br>EA | **rs6736741** | **2** | **144278534** | **C** | **0.180** | **1.02E-08** | **1.19** | **0.03** | **ARHGAP15:RP11-570L15.2:RP11-570L15.1** | **ncRNA_intronic** | **0.021** | **5** |
|  | rs112625468 | 17 | 76192318 | T | 0.443 | 5.15E-07 | 0.89 | 0.02 | AFMID | intronic | 0.272 | 7 |
|  | rs2452920 | 17 | 70949903 | C | 0.377 | 6.59E-07 | 0.89 | 0.02 | SLC39A11 | intronic | 3.624 | 5 |
| Diverticulitis | **rs4662208** | **2** | **144338448** | **A** | **0.177** | **3.22E-10** | **1.46** | **0.06** | **ARHGAP15:RP11-570L15.1** | **ncRNA_intronic** | **0.922** | **7** |
| EA | rs12671172 | 7 | 15227759<br>1 | G | 0.18<br>5 | 1.25E-07 | 0.67 | 0.0<br>8 | *AC104843.4* | intergenic | NA | 7 |
| Diverticulosis | rs11569231 | 3 | 14169238<br>1 | C | 0.12<br>7 | 1.74E-07 | 1.76 | 0.1<br>1 | *TFDP2* | intronic | 2.619 | NA |
| AA | rs114257184 | 8 | 12935767 | A | 0.14<br>5 | 8.56E-07 | 1.68 | 0.1<br>1 | *DLC1* | intergenic | 3.604 | 6 |
| Diverticulitis | rs7327483 | 13 | 85722171 | C | 0.29<br>7 | 8.33E-08 | 2.55 | 0.1<br>7 | *LINC00375* | intergenic | 3.128 | 4 |
| AA | rs140843945 | 5 | 12467136<br>2 | A | 0.02<br>8 | 1.11E-07 | 11.8<br>3 | 0.4<br>7 | *RN7SKP117* | intergenic | 4.134 | 7 |
|  | rs143460556 | 1 | 75625398 | G | 0.03<br>3 | 2.09E-07 | 5.73 | 0.3<br>4 | *LHX8* | intronic | 7.885 | 6 |
|  | rs80233487 | 18 | 34877201 | A | 0.02<br>3 | 2.17E-07 | 6.33 | 0.3<br>6 | *CELF4* | intronic | 2.813 | 5 |
|  | rs4657237 | 1 | 16288076<br>7 | C | 0.41<br>5 | 2.75E-07 | 2.46 | 0.1<br>8 | *C1orf110* | intergenic | 2.381 | 5 |
|  | rs6793498 | 3 | 39436360 | G | 0.18<br>1 | 2.96E-07 | 2.78 | 0.2<br>0 | *SLC25A38* | intronic | 2.778 | 7 |
|  | rs144422193 | 11 | 39890225 | G | 0.05<br>7 | 3.32E-07 | 3.93 | 0.2<br>7 | *RP11-810F22.1* | intergenic | 0.209 | 6 |
|  | rs78108838 | 10 | 11974029<br>2 | T | 0.07<br>0 | 4.29E-07 | 3.87 | 0.2<br>7 | *RAB11FIP2* | intergenic | 0.061 | 6 |
|  | rs74592875 | 8 | 58014100 | A | 0.04<br>4 | 5.79E-07 | 3.96 | 0.2<br>8 | *RNA5SP266* | intergenic | 3.428 | 7 |
|  | rs140818624 | 9 | 13611196<br>5 | A | 0.01<br>4 | 6.12E-07 | 10.5<br>3 | 0.4<br>7 | *LCN1P1* | intergenic | 0.563 | 5 |
|  | **rs14251961**<br>**7** | **5** | **64307317** | **G** | **0.01**<br>**2** | **6.19E-**<br>**07** | **12.2**<br>**4** | **0.5**<br>**0** | ***CWC27*** | **intronic** | **2.053** | **7** |
|  | rs4749487 | 10 | 30034377 | G | 0.05<br>1 | 6.95E-07 | 3.11 | 0.2<br>3 | *SVIL* | intergenic | 11.440 | 5 |
|  | rs11168732 | 12 | 49164120 | C | 0.45<br>5 | 7.81E-07 | 2.35 | 0.1<br>7 | *LINC00935, ADCY6,*<br>*MIR4701* | intronic | 1.338 | 5 |

Boldface type indicates the variants that meet the genomewide significance. (p-value < 5E-08).
\* deleteriousness score (CADD score<sup>43</sup>)
\*\*potential regulatory functions (RegulomeDB score<sup>44</sup>)

**Figure 2.**
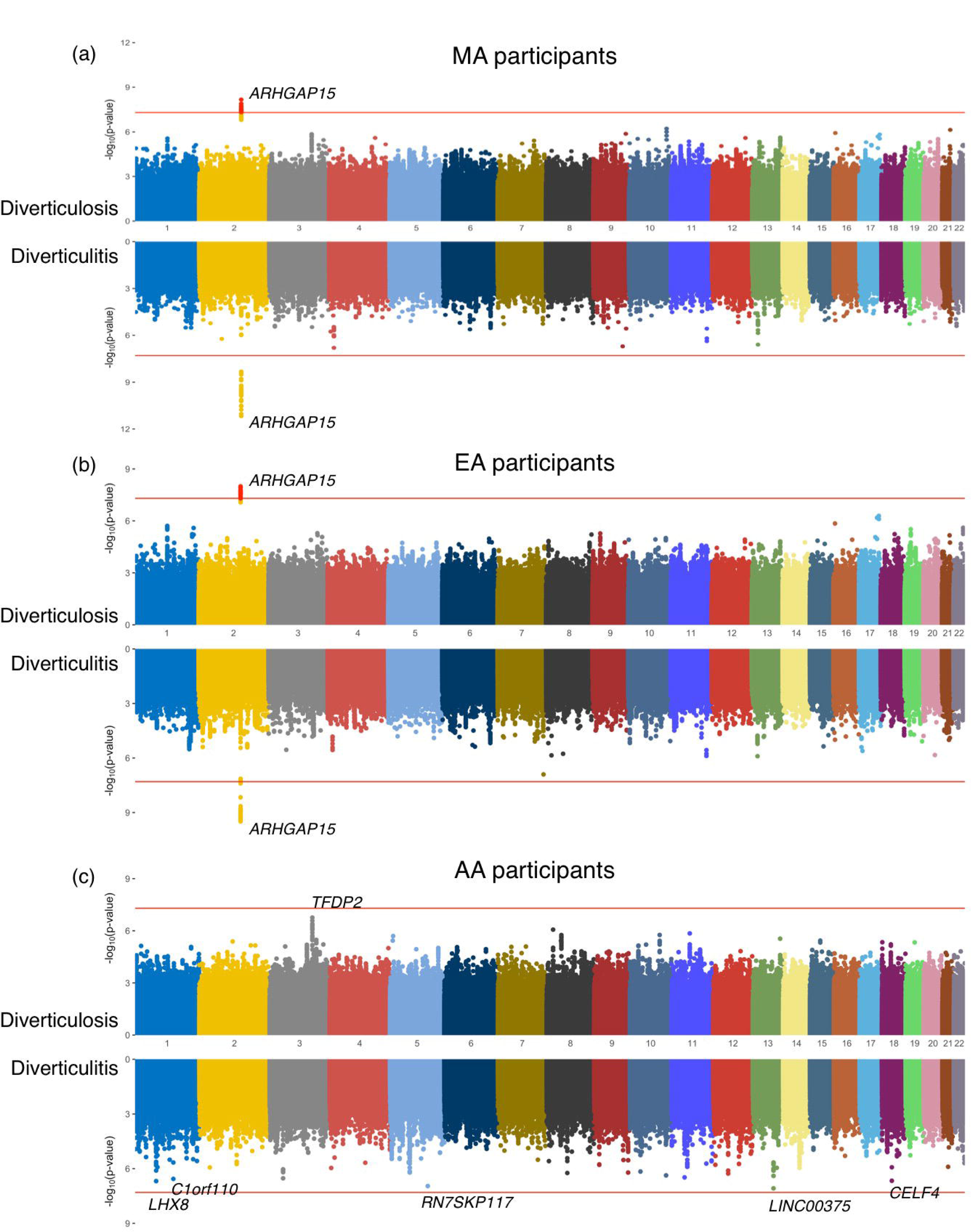
Manhattan plots of genome-wide associations with diverticular disease in (a) Multiancestry (MA) participants (n=21,777), (b) European Ancestry (EA) participants (n=19,211), and (c) African Ancestry (AA) participants (n=2,322). In each panel, the upper graph presents GWAS results of diverticulosis, and the bottom graph shows GWAS results of diverticulitis. The red horizontal line indicates genome-wide significance of p<5.0E-08 for each analysis.

The genetic signals found from EA-specific analysis and MA analysis were largely analogous, possibly due to the fact that approximately 85.0% of the discovery population was EA **(Figure 2, Table 1 and 3)**. Even though *ARHGAP15* loci showed non-significant p-values ranging from 0.24 up to 0.99 in the AA GWAS, the effect directions of *ARHGAP15* loci were largely positive and similar with EA GWAS of diverticular disease. The ORs of the variants were not as large as found in MA or EA results, but we confirmed the similar pattern of GWAS signals persisted especially intensified in diverticulitis GWAS in AA, ranging from 1.111 to 1.464, except a couple of them showed negative ORs (0.837 and 0.994) **(Supplementary Table 3)**.

We performed additional GWAS in the ICD-phenotyped diverticulitis cohort, and replicated a *ARHGAP15* locus on chromosome 2 (rs6717024) as genome-wide significant, similar to previous studies. Regardless of phenotyping algorithms, the impact of the *ARHGAP15* region on diverticular disease was found to be consistent. As the ICD-based phenotype cohort has a larger sample size than our NLP-enriched algorithm, it possibly identified a greater number of associations under the suggestive threshold (p-value< 1E-06). One of the strongest associations was rs11843418 (*FAM115A*), which was previously identified^23, 25^, but was not detected in our NLP-enriched GWAS possibly due to statistical power or different genetic composition of study cohorts. **(Supplementary Table 4)**

### LD score regression

A significant and positive genetic correlation was observed between diverticulosis and diverticulitis when comparing the effect sizes of those respective GWAS summary statistics: 0.935 (p-value=5.9E-03, SE=0.33) in MA, 0.997 (p-value=0.04, SE=0.48) in EA. The model does not converge in AA possibly due to low sample number. The ratio statistics of LDSC (0.68 and 0.35, for diverticulosis and diverticulitis respectively) reveals that polygenicity might not be the main driver of the observed signal in diverticular disease.

Calculated by LDSC, the genome-wide SNP base heritability was 4.2% (standard error (SE)=0.03) in diverticulosis and 23.7% (SE=0.15) in diverticulitis. LDSC intercept were stable and close to 1 (1.02 for both diverticulosis and diverticulitis), which means that the confounding factors such as population stratification were adequately controlled.

### PheWAS

#### (1) Diverticular disease susceptibility variants identified in our MA, EA, AA GWAS (p < 1E-06) tested in the medical phenome of MA, EA, AA participants

We observed FDR-significant PheWAS associations (FDR < 0.05) between diverticular disease phecodes (562, 562.1, and 562.2) and several independent (LD r^2^<0.1) *ARHGAP15* loci in MA and EA PheWAS **(Table 4)**. Other than diverticular EHR phenotypes, rs9565028 (*NBEA* gene) shows FDR-significant associations with genitourinary manifestations including ‘functional disorders of bladder’ (phecode 596.5) and ‘other disorders of bladder’ (phecode 596) in the MA and EA phenome. No significant associations were identified in AA PheWAS.

**Table 4:**
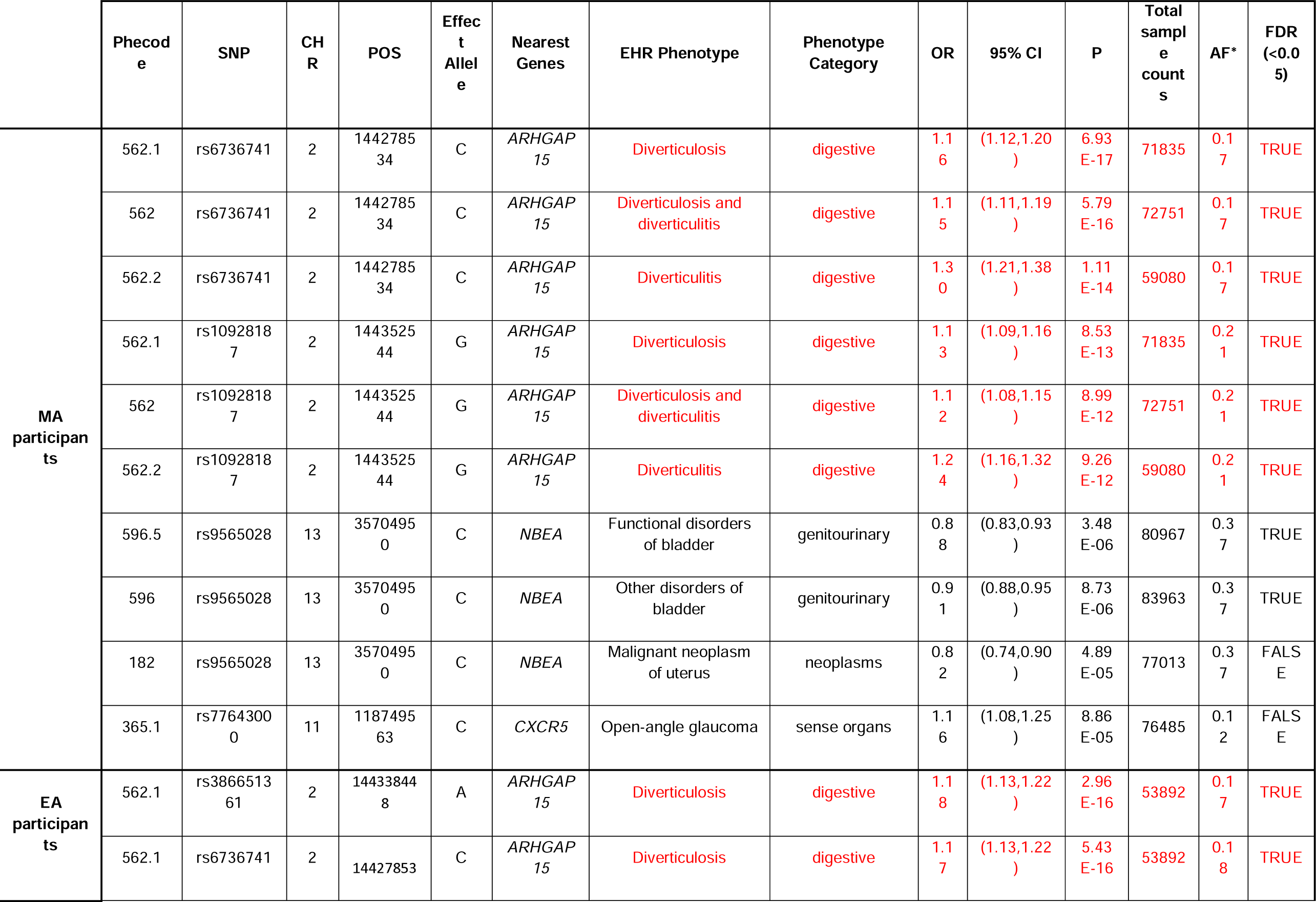
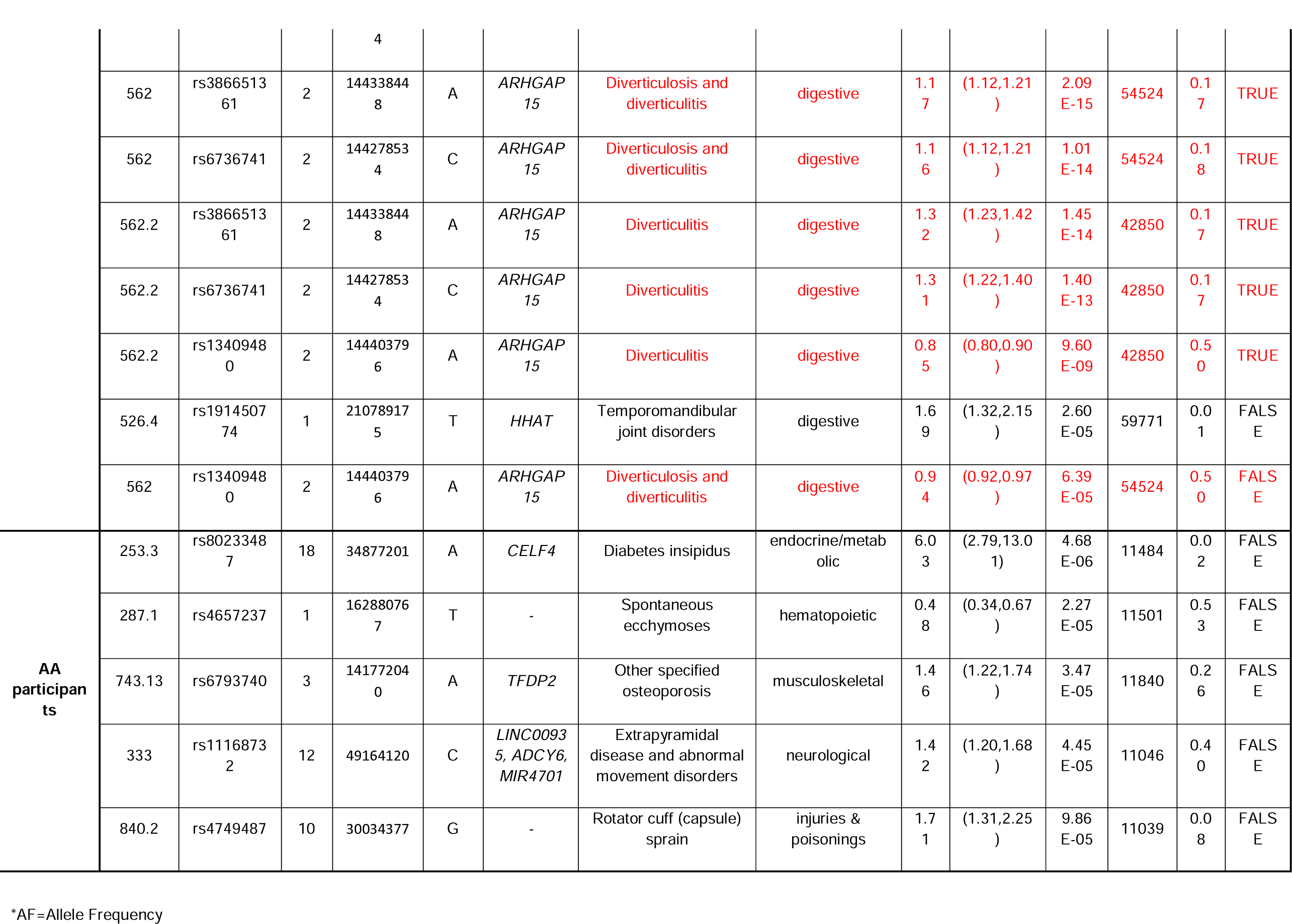
Significant genotype-EHR phenotype associations (suggestive threshold P<1E-04) from ancestry-stratified PheWAS of the discovered Diverticular disease susceptibility SNPs from our GWAS

#### (2) Diverticular disease susceptibility variants identified in previous GWAS (p < 5E-08) tested in the medical phenome of MA, EA, AA participants

In the MA PheWAS, 55 genotype-EHR phenotype associations confirmed FDR significance and reported in **Supplementary Table 1**. Among them, 18 significant genotype-EHR associations were endocrine/metabolic phenotypes, 17 of them were digestive phenotypes and 10 of them were circulatory system related phenotypes. The largest number of significant EHR phenotype associations were diverticular disease; 7 ‘diverticulosis and diverticulitis’, 7 ‘diverticulosis’ and 1 ‘diverticulitis’ were identified significant in MA PheWAS. Other than the *ARHGAP15* loci, rs4333882 (*SLC35F3* gene) and rs10472291 (*WDR70* gene) showed significant clinical associations with diverticular disease.

SNP rs9272785 (*HLA-DQA1* gene, proxy variant for rs7990) generated the most significant association in MA PheWAS coupled with ‘rheumatoid arthritis’ (phecode 714.1). The SNP was also strongly associated with several diabetes manifestations, including ‘type 1 diabetes’ (phecode 250.1), ‘type 1 diabetes with ophthalmic’ (phecode 250.13), ‘type 1 diabetes with ketoacidosis’ (phecode 250.11), ‘type 2 diabetes’ (phecode 250.2), etc.

In the EA PheWAS, 49 genotype-EHR phenotype associations were identified with FDR significance **(Supplementary table 1)**. Among them, 17 EHR phenotypes are classified as digestive phenotypes, 15 are endocrine/metabolic related phenotypes and 6 are related to circulatory system. Rs9272785 (*HLA-DQA1* gene) also marked the most significant association in EA PheWAS with ‘rheumatoid arthritis’ (phecode 714.1). The variant also revealed additional associations in EA phenome, including ‘developmental delays and disorders’ (phecode 315), ‘multiple sclerosis’ (phecode 335), ‘ulcerative colitis’ (phecode 555.2) and ‘chronic lymphocytic thyroiditis’ (phecode 256.21).

In AA PheWAS, two genotype-EHR phenotype associations met the FDR significance of 0.05: rs9272785 (*HLA-DQA1* gene) displayed the most significant SNP-phenotype association as it did in MA and EA PheWAS. The variant showed its strong associations with ‘type 1 diabetes with ketoacidosis’ (phecode 250.11) and ‘type 1 diabetes’ (phecode 250.1) in AA phenome.

## Discussion

To date, patient identification in previous GWAS studies were partially limited in that they mostly used inpatient medical coding which might resulted in under-diagnosis of the case patients as imaging reports were not used, and/or misclassification of controls who possibly have diverticular disease. In the most recent GWAS of diverticular disease^24^, the replication cohorts were manually reviewed with human hands-on input from physicians/technicians; however, manual review has a limited application to larger population-based datasets in its lack of scalability. Our phenotyping approach showed a significant improvement in performance (algorithm PPVs ≥ 0.94, 3.5 fold increase in diverticulosis patient identification) compared with use of only ICD-codes in patient classification, **(Table 2)** and supports the importance of leveraging the full breadth of data captured in EHRs^42, 43^.

Our transethnic GWASs of diverticular disease confirmed the strong genome-wide association of *ARHGAP15* in both diverticulosis and diverticulitis. **(Table 3)** *ARHGAP15* is known to strongly and negatively regulate GTPase binding property of the Rac protein family in leukocytes, which modulates important antimicrobial functions^44^. In an *in vivo* model of severe abdominal sepsis, null *ARHGAP15* knockout mice were reported to recruit more neutrophils to the site of infection; thereby limiting infection spread, systematic inflammation and bacterial growth^44^. This mechanism of *ARHGAP15* possibly impacts the inflammatory environment of the intestine, promoting the development of diverticula or progression of diverticula due to bacterial growth along the colonic wall.

Ancestry-stratified GWAS analyses revealed that the often-replicated associations between *ARHGAP15* with diverticular disease in EA cohorts, and similar positive effect sizes but little to no association observed in AA cohorts **(Supplementary Table 3)**. It is of note that the sample size for the AA cohort is less than 1/10^th^ that of the EA cohort, as well as different risk allele frequencies between ancestries. Our additional power calculation showed that at least 15,000 participants are needed to perform GWAS in *ARHGAP15* loci (EAF 0.18, disease prevalence 0.10, OR 1.20) with 80% statistical power **(Supplementary Figure 1)**. Further investigation is needed to confirm the universal susceptibility effect of *ARHGAP15* to diverticular disease in non-European ancestry.

Our PheWAS assessed the clinical associations of the GWAS variants of diverticular disease with EHR-phenotypes **(Table 4)**. PheWAS of the independent *ARHGAP15* loci (rs6736741, rs10928187, rs386651361) confirms its significant phenotypic expression with diverticular disease in MA and EA along with the second most significant association of ‘paralytic ileus’. Some genitourinary phenotypes of functional bladder disorders found in MA and EA should be noted in that the muscular motility or neuromuscular dysfunction of internal organs possibly influence both colonic wall for diverticulosis and bladder muscle for urinary disorder.

In the PheWAS of the established diverticular variants, we identified several circulatory system related EHR phenotypes associated with diverticular disease variants, including phlebitis and thrombophlebitis (phecode 451), pulmonary heart disease (phecode 415, 415.1), and deep vein thrombosis (phecode 452.2). Interestingly, recent studies have suggested a possible epidemiologic association between diverticular disease and acute coronary syndromes and thromboembolic events^45, 46^. We also confirmed the associations of rs9272785 (*HLA-DQA1* gene) with type 1 diabetes and its manifestations with FDR significance across ancestries. The HLA class 2 region, where rs9272785 is located, is not only associated with risk of type 1 diabetes but also increased susceptibility for juvenile rheumatoid arthritis and other autoimmune diseases^47, 48^.

The relational investigation with LD score regression demonstrates substantial genetic overlap between diverticulosis and diverticulitis, as high as 0.935 (correlation p-value=5.9E-03, SE=0.33) in MA, 0.997 (correlation p-value=0.04, SE=0.48) in EA. In addition, we observed the intensified GWAS signals in diverticulitis patients over diverticulosis cases throughout our GWAS analyses. The high genetic similarity between both traits is somewhat expected, since diverticulitis cannot be developed in the absence of diverticulosis.

Compared to previous diverticular GWASs, our summary statistics generally show larger effect sizes possibly fueled by the improved accuracy of our colonoscopy and abdominal imaging-based phenotyping algorithm. For example, rs6734367, the strongest *ARHGAP15* locus reported in Maguire et al^23^ showed positive OR of 1.010 in the original study, whereas it presents a stronger OR as high as 1.177 (diverticulosis) and 1.280 (diverticulitis) in our EA GWAS with the same allelic direction **(Supplementary Table 6)**. For the rest of the genome-wide significant SNPs identified by Maguire et al^23^, the ORs in our study overwhelmingly show increased effect sizes despite a cohort 1/20^th^ the size of Maguire et al. **(Figure 3)**. As cohort size gets larger and diverse patients with different genetic background are included, our results suggest the improved analytical power for future genomic research with the integration of different layers of the EHR data.

**Figure 3.**
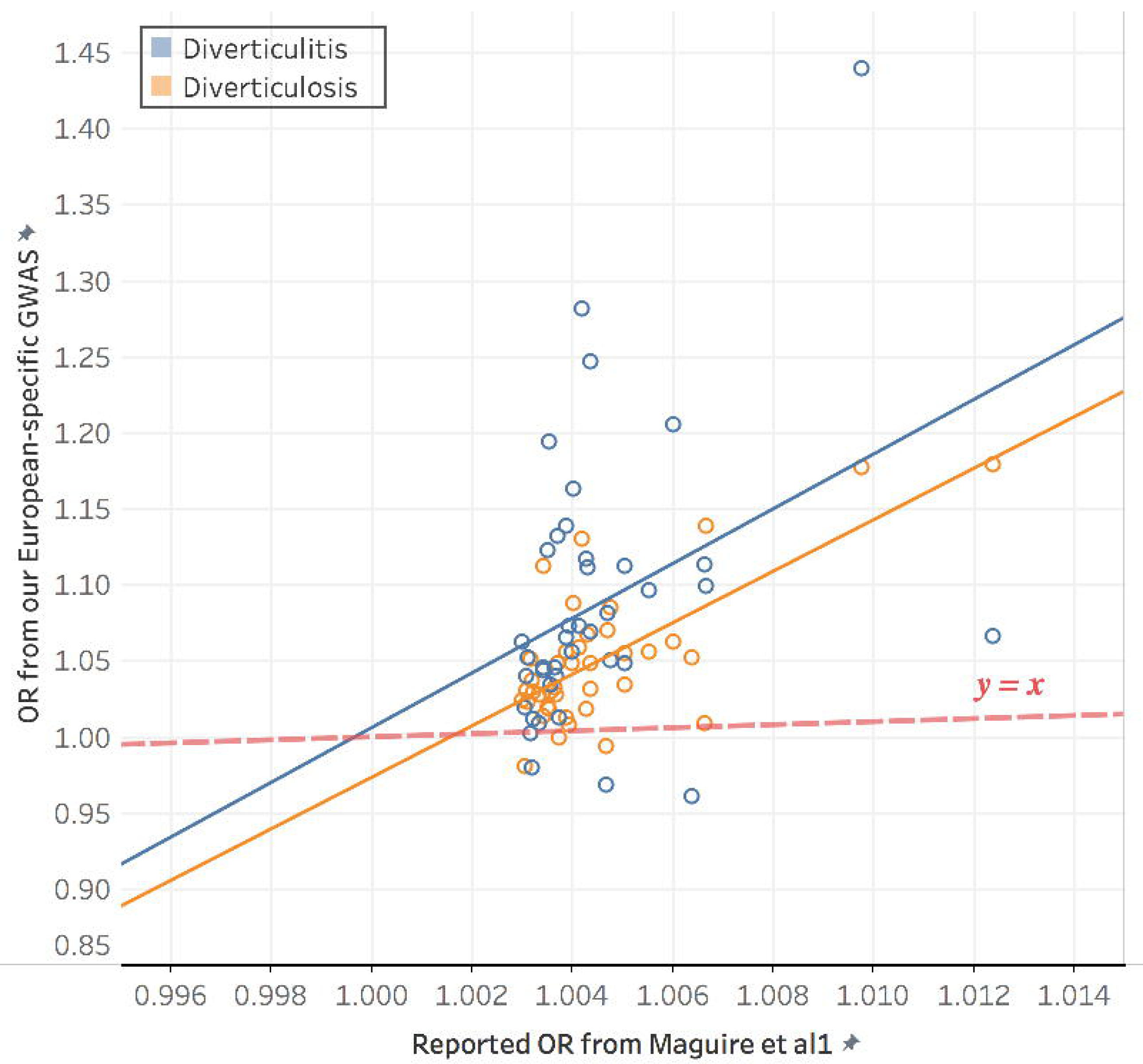
Comparison of effect size (OR) between our GWAS with NLP-enriched phenotyping and previous GWAS with ICD-based phenotyping from Maguire et al. The dashed y=x line indicates equal ORs in both studies.

There are several caveats in interpreting our findings. We did not separately validate our phenotyping algorithms’ performance for diverticulitis vs. diverticulosis, which should be done in future research. Possibly due to reduced statistical power with multiple testing, our GWAS did not identify any novel association and only confirmed an existing locus with diverticular disease, albeit with the larger effect sizes across the analyses. Also, Our MA analysis is composed of 85% EA participants, so the signals are largely driven by EA-centric results. The cohort size of AA is considerably smaller than EA or MA cohort, which elevates the risk for false positive findings.

In summary, our multi-institutional EHR study of genotype-phenotype associations in diverticular disease performed sectional analysis of diverticulosis and diverticulitis among the different genetic races, which revealed the possible transethnic genetic effect of *ARHGAP15* loci with diverticular disease across European and African ancestries. By implementing NLP, our approach showcased a comprehensive use of different formats of EHR data from GWAS to post-GWAS functional analysis including PheWAS. The enhanced methods have shown the efficiency and feasibility of EHR information in disease case/control classification with relevant informatics techniques, and a clinical possibility for an integrative analytical pipeline to facilitate etiological investigation of a disease.

## Abbreviations

AA: African ancestry;
ADPKD: autosomal dominant polycystic kidney disease;
BMI: body mass index;
CI: Confidence interval;
EA: European ancestry;
EHR: electronic health record;
eMERGE: Electronic Medical Records and Genomics network;
FDR: false discovery rate;
GI: gastrointestinal;
GWAS: genome-wide association study;
HWE: Hardy-Weinberg equilibrium;
KPWA/UW: Kaiser Permanente Washington/ University of Washington;
LD: linage disequilibrium;
MAF: minor allele frequency;
NLP: natural language processing;
NU: Northwestern University;
PCA: Principal Components Analysis;
PheWAS: phenomewide association study;
PPV: positive predictive value;
EAF: effect allele frequency;
SNP: single nucleotide variant;
VU: Vanderbilt University

---

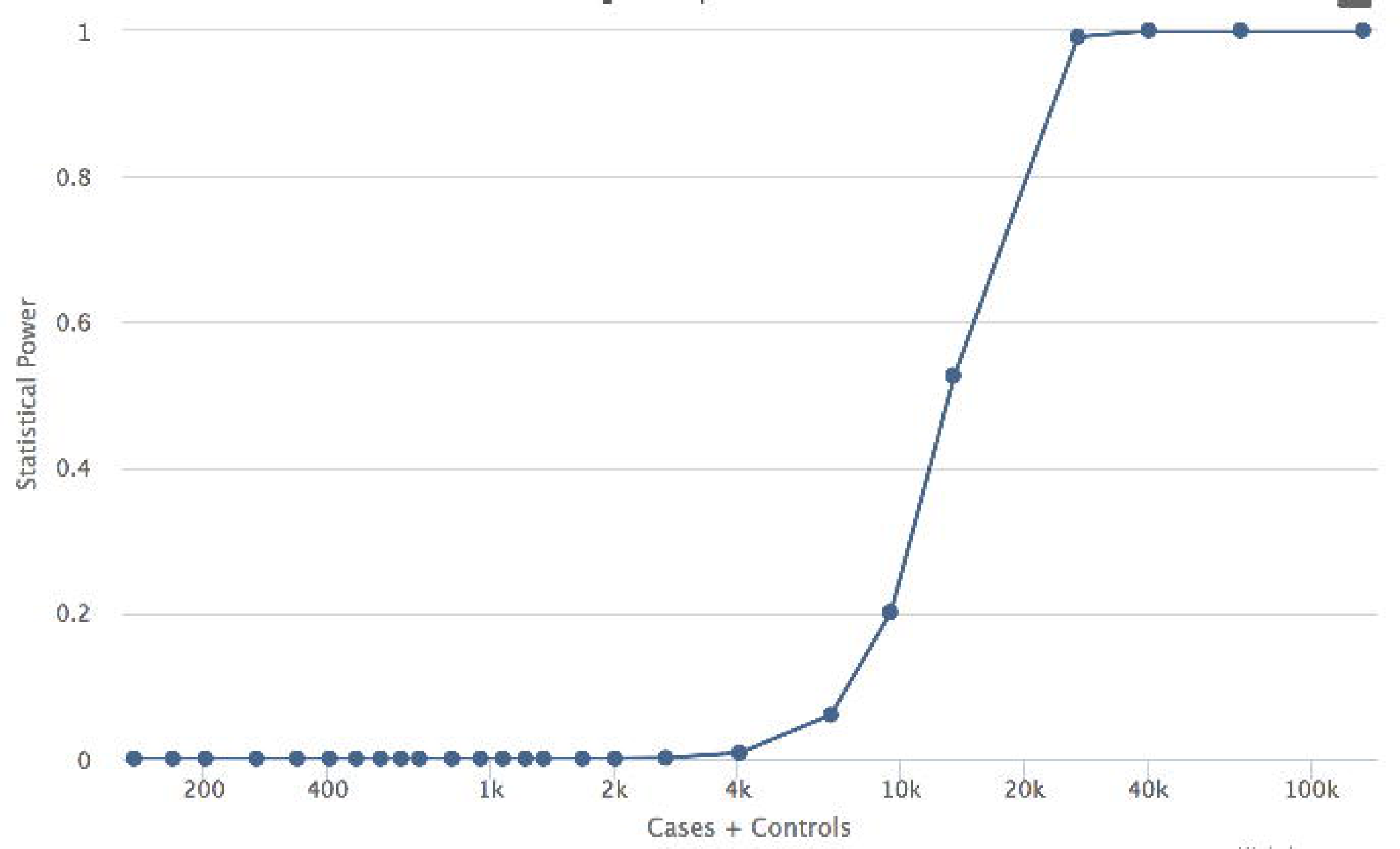

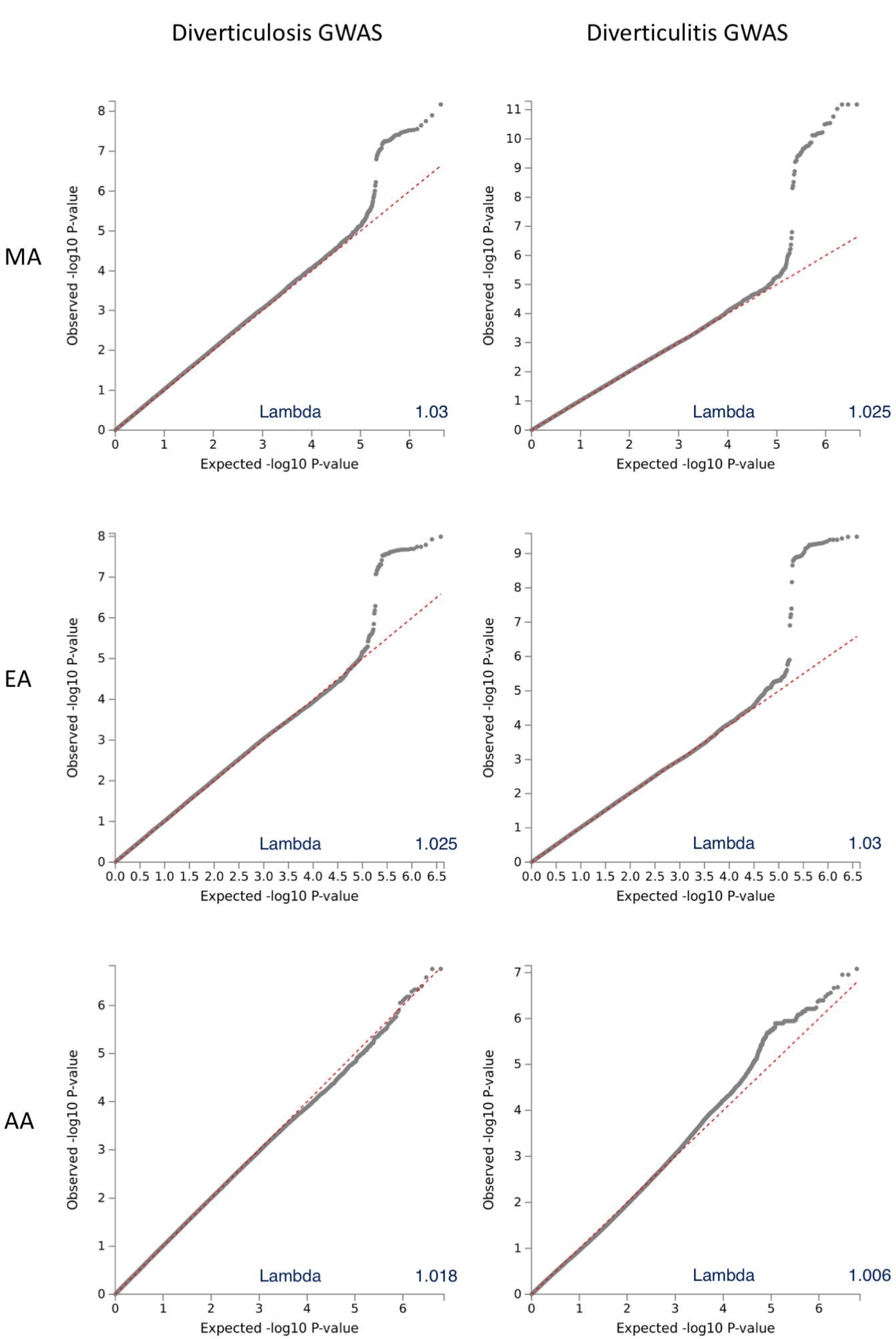

## References

1. Sandler RS, Everhart JE, Donowitz M, et al. The burden of selected digestive diseases in the United States. Gastroenterology 2002;122:1500–11.

2. Peery AF, Crockett SD, Barritt AS, et al. Burden of Gastrointestinal, Liver, and Pancreatic Diseases in the United States. Gastroenterology 2015;149:1731–1741 e3.

3. Peery AF, Crockett SD, Murphy CC, et al. Burden and Cost of Gastrointestinal, Liver, and Pancreatic Diseases in the United States: Update 2018. Gastroenterology 2019;156:254–272 e11.

4. Strate LL, Morris AM. Epidemiology, Pathophysiology, and Treatment of Diverticulitis. Gastroenterology 2019;156:1282–1298 e1.

5. Patel. GSALAFK. Diverticulosis. StatPearls [Internet]. 2019 Apr 11 ed, 2019.

6. Reichert MC, Lammert F. The genetic epidemiology of diverticulosis and diverticular disease: Emerging evidence. United European Gastroenterol J 2015;3:409–18.

7. Colcock BP. Diverticular disease of the colon. Major Probl Clin Surg 1971;11:1–135.

8. Shahedi K, Fuller G, Bolus R, et al. Long-term risk of acute diverticulitis among patients with incidental diverticulosis found during colonoscopy. Clin Gastroenterol Hepatol 2013;11:1609–13.

9. Painter NS, Burkitt DP. Diverticular disease of the colon: a deficiency disease of Western civilization. Br Med J 1971;2:450–4.

10. Painter NS, Burkitt DP. Diverticular disease of the colon, a 20th century problem. Clin Gastroenterol 1975;4:3–21.

11. Makela J, Kiviniemi H, Laitinen S. Prevalence of perforated sigmoid diverticulitis is increasing. Dis Colon Rectum 2002;45:955–61.

12. Etzioni DA, Mack TM, Beart RW, Jr., et al. Diverticulitis in the United States: 1998-2005: changing patterns of disease and treatment. Ann Surg 2009;249:210–7.

13. Nagata N, Niikura R, Aoki T, et al. Increase in colonic diverticulosis and diverticular hemorrhage in an aging society: lessons from a 9-year colonoscopic study of 28,192 patients in Japan. Int J Colorectal Dis 2014;29:379–85.

14. Warner E, Crighton EJ, Moineddin R, et al. Fourteen-year study of hospital admissions for diverticular disease in Ontario. Can J Gastroenterol 2007;21:97–9.

15. Kang JY, Hoare J, Tinto A, et al. Diverticular disease of the colon--on the rise: a study of hospital admissions in England between 1989/1990 and 1999/2000. Aliment Pharmacol Ther 2003;17:1189–95.

16. Lee YS. Diverticular disease of the large bowel in Singapore. An autopsy survey. Dis Colon Rectum 1986;29:330–5.

17. Ogunbiyi OA. Diverticular disease of the colon in Ibadan, Nigeria. Afr J Med Med Sci 1989;18:241–4.

18. Walker AR, Segal I. Epidemiology of noninfective intestinal diseases in various ethnic groups in South Africa. Isr J Med Sci 1979;15:309–13.

19. Aldoori WH, Giovannucci EL, Rimm EB, et al. A prospective study of diet and the risk of symptomatic diverticular disease in men. Am J Clin Nutr 1994;60:757–64.

20. Aldoori WH. The protective role of dietary fiber in diverticular disease. Adv Exp Med Biol 1997;427:291–308.

21. Peery AF, Barrett PR, Park D, et al. A high-fiber diet does not protect against asymptomatic diverticulosis. Gastroenterology 2012;142:266–72 e1.

22. Peery AF, Sandler RS, Ahnen DJ, et al. Constipation and a low-fiber diet are not associated with diverticulosis. Clin Gastroenterol Hepatol 2013;11:1622–7.

23. Maguire LH, Handelman SK, De X, et al. Genome-wide association analyses identify 39 new susceptibility loci for diverticular disease. Nat Genet 2018;50:1359–1365.

24. Schafmayer C, Harrison JW, Buch S, et al. Genome-wide association analysis of diverticular disease points towards neuromuscular, connective tissue and epithelial pathomechanisms. Gut 2019;68:854–865.

25. Sigurdsson S, Alexandersson KF, Sulem P, et al. Sequence variants in ARHGAP15, COLQ and FAM155A associate with diverticular disease and diverticulitis. Nat Commun 2017;8:15789.

26. Matrana MR, Margolin DA. Epidemiology and pathophysiology of diverticular disease. Clin Colon Rectal Surg 2009;22:141–6.

27. Destigter KK, Keating DP. Imaging update: acute colonic diverticulitis. Clin Colon Rectal Surg 2009;22:147–55.

28. Feingold D, Steele SR, Lee S, et al. Practice parameters for the treatment of sigmoid diverticulitis. Dis Colon Rectum 2014;57:284–94.

29. Joseph DA KJ, Richards TB, Thomas CC, Richardson LC. Use of colorectal cancer screening tests by state. Preventing Chronic Disease 2018;15:170535.

30. Denny JC, Crawford DC, Ritchie MD, et al. Variants near FOXE1 are associated with hypothyroidism and other thyroid conditions: using electronic medical records for genome- and phenome-wide studies. Am J Hum Genet 2011;89:529–42.

31. Stanaway IB, Hall TO, Rosenthal EA, et al. The eMERGE genotype set of 83,717 subjects imputed to ∼40 million variants genome wide and association with the herpes zoster medical record phenotype. Genet Epidemiol 2019;43:63–81.

32. Newton KM, Peissig PL, Kho AN, et al. Validation of electronic medical record-based phenotyping algorithms: results and lessons learned from the eMERGE network. J Am Med Inform Assoc 2013;20:e147–54.

33. Chang CC, Chow CC, Tellier LC, et al. Second-generation PLINK: rising to the challenge of larger and richer datasets. Gigascience 2015;4:7.

34. Kircher M, Witten DM, Jain P, et al. A general framework for estimating the relative pathogenicity of human genetic variants. Nat Genet 2014;46:310–5.

35. Boyle AP, Hong EL, Hariharan M, et al. Annotation of functional variation in personal genomes using RegulomeDB. Genome Res 2012;22:1790–7.

36. Yang J, Lee SH, Goddard ME, et al. GCTA: a tool for genome-wide complex trait analysis. Am J Hum Genet 2011;88:76–82.

37. Denny JC, Bastarache L, Ritchie MD, et al. Systematic comparison of phenome-wide association study of electronic medical record data and genome-wide association study data. Nat Biotechnol 2013;31:1102–10.

38. Bulik-Sullivan B, Finucane HK, Anttila V, et al. An atlas of genetic correlations across human diseases and traits. Nat Genet 2015;47:1236–41.

39. Bulik-Sullivan BK, Loh PR, Finucane HK, et al. LD Score regression distinguishes confounding from polygenicity in genome-wide association studies. Nat Genet 2015;47:291–5.

40. Carroll RJ, Bastarache L, Denny JC. R PheWAS: data analysis and plotting tools for phenome-wide association studies in the R environment. Bioinformatics 2014;30:2375–6.

41. Erichsen R, Strate L, Sorensen HT, et al. Positive predictive values of the International Classification of Disease, 10th edition diagnoses codes for diverticular disease in the Danish National Registry of Patients. Clin Exp Gastroenterol 2010;3:139–42.

42. Wei WQ, Denny JC. Extracting research-quality phenotypes from electronic health records to support precision medicine. Genome Med 2015;7:41.

43. Peissig PL, Rasmussen LV, Berg RL, et al. Importance of multi-modal approaches to effectively identify cataract cases from electronic health records. J Am Med Inform Assoc 2012;19:225–34.

44. Costa C, Germena G, Martin-Conte EL, et al. The RacGAP ArhGAP15 is a master negative regulator of neutrophil functions. Blood 2011;118:1099–108.

45. Pers TH, Karjalainen JM, Chan Y, et al. Biological interpretation of genome-wide association studies using predicted gene functions. Nat Commun 2015;6:5890.

46. Strate LL, Erichsen R, Horvath-Puho E, et al. Diverticular disease is associated with increased risk of subsequent arterial and venous thromboembolic events. Clin Gastroenterol Hepatol 2014;12:1695–701 e1.

47. Begovich AB, Bugawan TL, Nepom BS, et al. A specific HLA-DP beta allele is associated with pauciarticular juvenile rheumatoid arthritis but not adult rheumatoid arthritis. Proc Natl Acad Sci U S A 1989;86:9489–93.

48. Noble JA, Valdes AM. Genetics of the HLA region in the prediction of type 1 diabetes. Curr Diab Rep 2011;11:533–42.

